# Best practices for single-cell histone modification analysis

**DOI:** 10.1101/2022.09.21.508811

**Authors:** Félix Raimundo, Pacôme Prompsy, Jean-Philippe Vert, Céline Vallot

## Abstract

**Background:** Single-cell histone post translation modification (scHPTM) assays such as scCUT&Tag or scChIP-seq allow single-cell mapping of diverse epigenomic landscapes within complex tissues, and are likely to unlock our understanding of various epigenetic mechanisms involved in development or diseases. Running an scHTPM experiment and analyzing the data produced remains, however, a challenging task since few consensus guidelines exist currently regarding good practices for experimental design and data analysis pipelines.

**Methods:** We perform a computational benchmark to assess the impact of experimental parameters and of the data analysis pipeline on the ability of the cell representation produced to recapitulate known biological similarities. We run more than ten thousands experiments to systematically study the impact of coverage and number of cells, of the count matrix construction method, of feature selection and normalization, and of the dimension reduction algorithm used.

**Results:** The analysis of the benchmark results allows us to identify key experimental parameters and computational choices to obtain a good representation of single-cell HPTM data. We show in particular that the count matrix construction step has a strong influence on the quality of the representation, and that using fixed-size bin counts outperforms annotation-based binning; that dimension reduction methods based on latent semantic indexing outperform others; and that feature selection is detrimental, while keeping only high-quality cells has little influence on the final representation as long as enough cells are analyzed.

## Introduction

Posttranslational modifications (PTM) of histone proteins are key epigenetic events that modulate chromatin structure, nucleosome positioning and transcription. They are involved in numerous biological processes, including DNA repair [1], development [2, 3] and cancer [4]. With the recent advent of high-throughput technologies to measure histone PTM at the single-cell level (scHPTM), such as single-cell chromatin immunoprecipitation followed by sequencing (scChIP-seq) [5] and single-cell cleavage under targets and tagmentation (scCUT&Tag) [6, 7], it is now feasible to explore the diversity of histone PTM in complex biological samples with an ever-increasing level of details [8, 6, 9, 10]. ScHPTM has already allowed new biological insights such as the discovery of epigenetic factors involved in cancer response to chemotherapy [11], and is likely to unlock our understanding of various epigenetic mechanisms in the years to come.

While scHPTM has great potential, it is also a relatively recent approach which comes with numerous computational challenges that need to be addressed in order to fully deliver its promise of capturing biologically relevant information from raw experimental data. In this work, we leave aside the question of which technology to use to generate scHPTM data, and focus instead on two important questions for practitioners, namely, 1) how to design experiments, in particular to choose a good trade-off between number of cells and coverage, and 2) how to computationally analyze the raw experimental data and transform them in biologically relevant representations, where subsequent analysis such as cell classification or lineage inference become feasible. While both questions have been investigated through systematic benchmarks and comparisons for more mature single-cell technologies such as single-cell RNA-seq (scRNA-seq) and single-cell sequencing assay for transposase-accessible chromatin (scATAC-seq) [12, 13, 14, 15, 16], we are not aware of any similar study conducted for the burgeoning field of scHPTM, leaving experimentalists without rational guidelines on how to design their scHPTM experiments and analyze the data they produce.

Given the similar nature of raw experimental data between scHPTM and scATAC-seq, namely, sequencing reads capturing an epigenomic signal distributed in specific regions over the whole genome, it would seem natural to use the same computational methods to analyze scHPTM and scATAC-seq data. However, both modalities differ in many aspects. First, the actual distribution of reads can be drastically different between scHPTM and ATAC-seq. Indeed ATAC-seq reads are known to cluster in relatively small, ∼1k base pairs (kbp), regions [17], whereas the regulatory regions for scHPTM vary much more widely in size (e.g., between 5kbp and 2000kbp for H3K27me3 [17]) and their locations can vary depending on the histone mark - from enhancers (H3K27ac) to gene body (H3K36me3) or intergenic regions (H3K27me3). Second, with current technologies, the number of sequenced reads in scHPTM is generally between a few hundred and a few thousand per cells, compared to several thousands for scATAC-seq and tens of thousands for scRNA-seq. Such a low coverage leads to only about 1% of the expected enriched regions to contain at least one read per cell (compared to 1-10% for scATAC-seq and 10-45% for scRNA-seq [12]). Thus one can not assume that computational recommendations for scATAC-seq or RNA-seq hold for scHPTM.

To start filling this gap, we perform in this paper a large-scale computational study to evaluate the impact and best choices for the number of cells, coverage per cell, cell selection, matrix construction algorithm, feature selection and dimension reduction algorithm. To quantify the impact of each of these factors, we use two single-cell multi-omics datasets where, in addition to scHPTM, a second modality is measured for each cell (gene expression or cell surface proteins); we then assess how well the cell-to-cell similarity observed with scHPTM data analysis agrees with the one inferred from the co-assay (RNA or protein) [18, 19]. The analysis of more than 10.000 computational experiments allows us to clarify the impact of various experimental choices and data processing factors for scHTPM data, and to suggest practical guidelines.

## Results

### Benchmarking methods for scHPTM analysis

Irrespective of the technology used, most protocols for scHPTM analysis produce sequencing reads which, after being mapped to a reference genome, indicate where on the genome a given PTM mark is likely to be present in each individual cell under study. A number of computational steps are then applied to transform these raw data into a useful representation of each individual cell, where downstream applications such as cell classification or differential analysis are performed. Here we focus on computational frameworks that produce a representation of each cell as a vector of moderate dimension (typically, 10 to 50 dimensions), which has been found to be a powerful approach for scRNA-seq data analysis [20] and is currently the *de facto* standard for scATAC-seq and scHPTM as well [20]. Going from the mapped read to a vector representation for each cell involves a number a steps that we investigate in this study (Figure 1A), including 1) the binning of the mapped reads into genomic regions in order to create a cell *×* region count matrix to summarize the raw data, 2) various quality control (QC) preprocessing operations to filter out low-quality cells and regions, and 3) an embedding method to build the representation of each cell from the preprocessed count matrix. Each step can be performed in many different ways, and we propose a benchmark to assess the impact of each choice at each step on the final cell representation (Figure 1).

**Figure 1:**
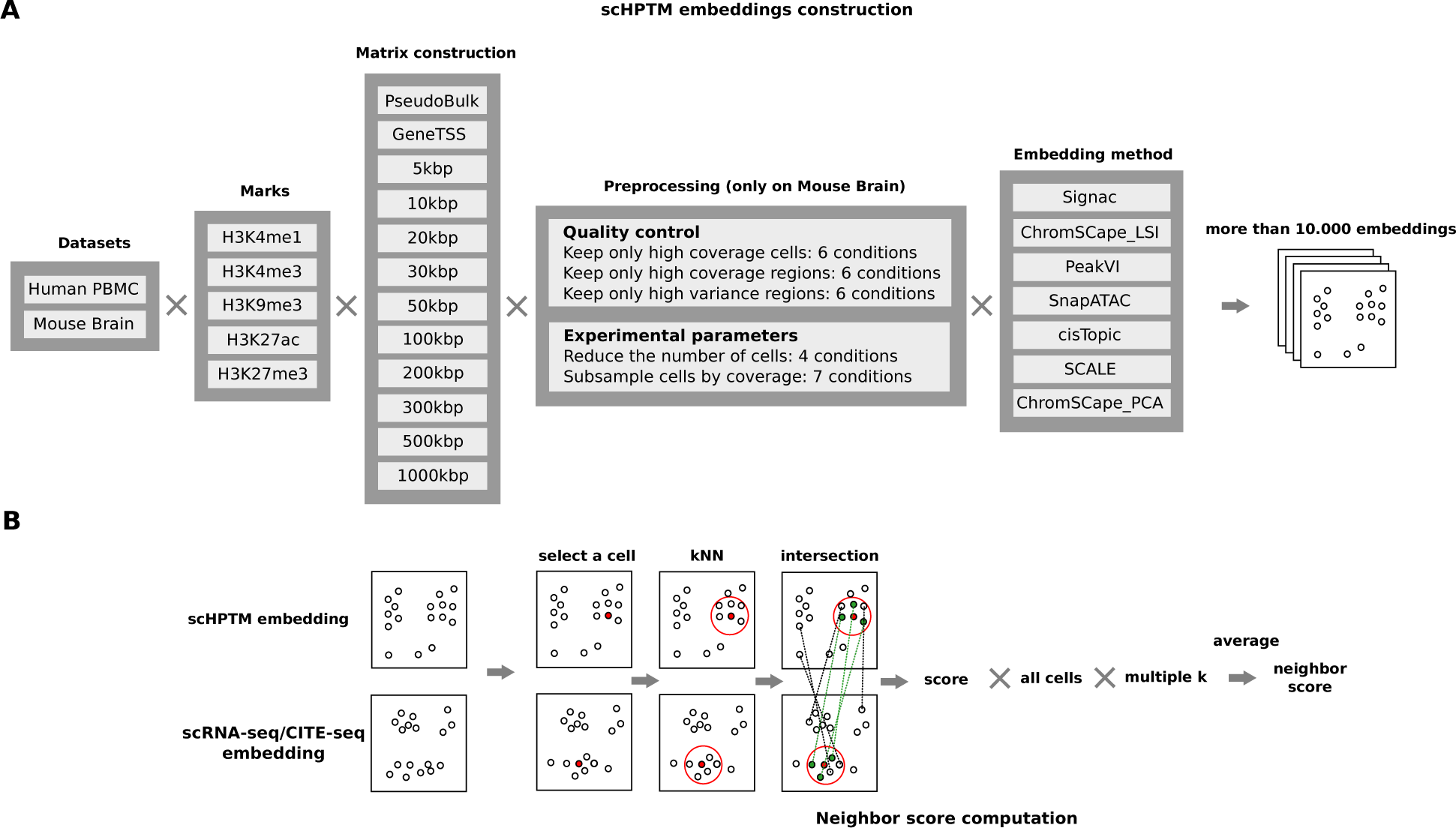
Overview of the evaluation protocol. **A**. We build the count matrix using different bin sizes as well as a GeneTSS annotation and peaks called on the pseudo bulk (only for the human PBMC dataset). We then simulate in-silico different experimental conditions for studying the role of the number of cells in a dataset, and the effect of the coverage per cell, as well as different feature selection strategies. Afterwards we run 7 different dimension reduction methods to obtain the cell representations. **B**. In order to compute the neighbor score, we start by selecting a cell, we then build the *k*NN graph for a value of *k* (5 in the figure), we then compute the size of the intersection between the neighborhood of the cell in the two embeddings (3 cells in the figure) and divide it by *k* to obtain the score for one cell and one value of k (score of 0.6 in the figure). We then compute and average this score over all the cells, to have an neighbor score for a given value of *k*, that score is then further averaged over different values of *k* (0.1%, 0.3%, 0.5%, 1%, 3%, 5% and 10% of the number of cells in the experience) to obtain the final neighbor score

In order to evaluate the impact of each decision on the quality of the final representation, we need reference datasets and a way to quantify the quality of that representation. As no ground truth reference datasets are available for scHPTM analysis, we rely on two datasets produced with multiomics co-assays (Table 1), where two modalities are measured simultaneously in each cell. More precisely, we consider a mouse brain dataset from [9] where five histone marks (H3K4m1, H3K4me3, H3K9me3, H3K27ac, and H3K27me3) are assessed by scHPTM jointly with scRNA-seq-based gene expression, and a human peripheral blood mononuclear cell (PBMC) dataset from [10] where the same five histone marks are assessed by scHPTM jointly with CITE-seq-based cell surface proteins. For both datasets, we use a unique representation of the second modality (respectively, scRNA-seq and CITE-seq) using a well-established method as a reference (scanpy’s [21] implementation of PCA), and compare each representation obtained from the scHPTM data to that reference. We compute a neighbor score that assesses to what extent neighbor cells in the scHPTM representation are also found neighbors in the reference representation of the second modality. The neighbor score varies between 0 when both representations disagree completely to 1 when both representations are identical (see Methods and Figure 1B). This evaluation has been previously used in [18, 19] and is currently the standard for evaluating modality alignment tasks in recent community benchmarks such as https://openproblems.bio/.

**Table 1:**
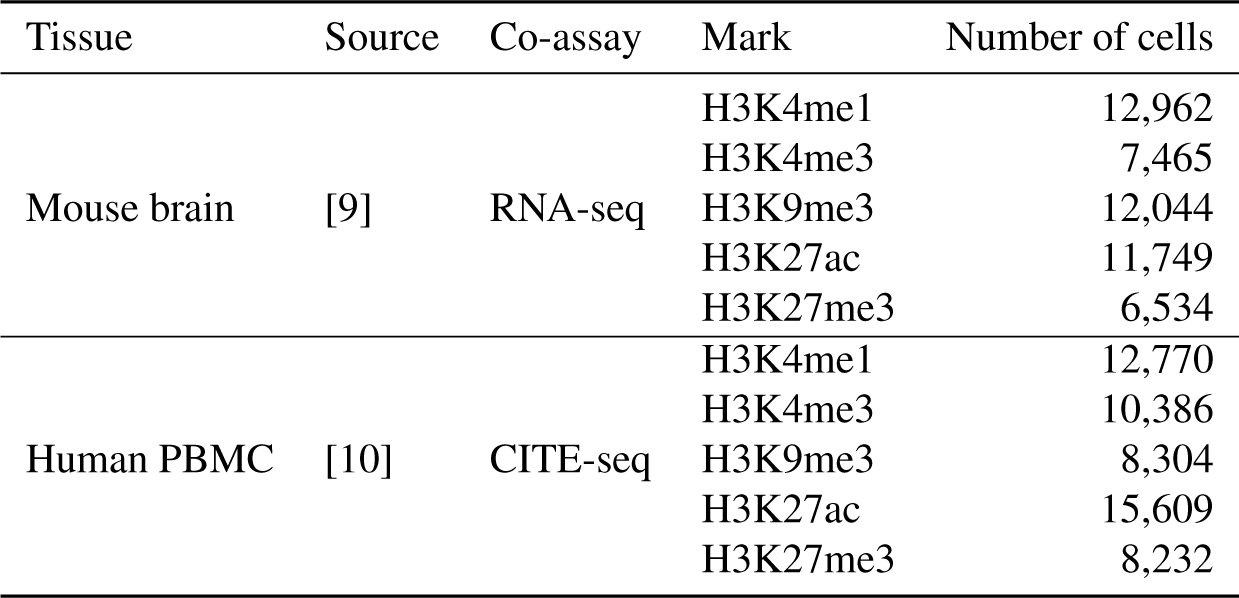
Description of the co-assay datasets used for this study

| Tissue | Source | Co-assay | Mark | Number of cells |
| --- | --- | --- | --- | --- |
| Mouse brain | [9] | RNA-seq | H3K4me1 | 12,962 |
|  |  |  | H3K4me3 | 7,465 |
|  |  |  | H3K9me3 | 12,044 |
|  |  |  | H3K27ac | 11,749 |
|  |  |  | H3K27me3 | 6,534 |
| Human PBMC | [10] | CITE-seq | H3K4me1 | 12,770 |
|  |  |  | H3K4me3 | 10,386 |
|  |  |  | H3K9me3 | 8,304 |
|  |  |  | H3K27ac | 15,609 |
|  |  |  | H3K27me3 | 8,232 |

For each dataset and each histone PTM mark, we systematically vary the choices that we can make in each step of the computational pipeline that goes from the mapped reads to the scHPTM representation of each cell, and measure the quality of the final representation with the neighbor score to assess the impact of the choices.

More precisely, for the first step that bins mapped reads to regions in order to build a first cell *×* region count matrix, we consider three different strategies that represent the various approaches used in practice for the analysis of single-cell epigenomic assays: 1) discretizing the whole genome into “bins” of fixed size, and trying different sizes following a logarithmic progression between 5kbp and 1000kbp, 2) counting the reads into bins based on prior biological knowledge, i.e., on genes and transcription start sites annotations (GeneTSS), 3) counting the reads into a set of peaks, characteristic of each cell population found in the sample (identified from the corresponding pseudo bulk using MACS2, [22] ‘PseudoBulk’). This last approach was only performed with the human PBMC dataset that is distributed in a format that allows us to build the pseudo bulk and use it for peak calling. With these matrices, we attempt different feature selection approaches to select only a subset of genomic regions to keep for further analysis: 1) selection of highly variable regions using Seurat’s [23] FindVariableFeatures function (variable features), 2) selection of regions with the highest coverage (top features). The first feature selection method is the current standard in scRNA-seq, and the second approach is recommended in Signac [24] for analyzing scATAC-seq. We further study the role of cell filtering based on their coverage, which is part of the standard analysis steps. For region filtering, we study the effect of coverage and variance filtering. We also simulate different experimental conditions *in silico* in order to evaluate how cell numbers affect cell representation, as well as the importance of their coverage. Finally, we consider seven popular methods for analyzing the count matrices: cisTopic [25], Signac [24] SnapATAC [26], PeakVI [18], SCALE [27], and ChromSCape [28] with TF-IDF (ChromSCape_LSI) and count per million (CPM) normalization (ChromSCape_PCA).

This leads us to test 11,970 combinations of mark, dimension reduction method, matrix construction, cell selection, feature selection, number of cells and coverage conditions, out of which 11,080 ran successfully (Tables S1-S2). Failures to run were generally due to memory issues on small bin sizes and GeneTSS annotation. We then analyze the impact of each decision choice and experimental condition by assessing statistically how the neighbor score of the representation varies with the decision.

### LSI based methods outperform other methods

We first focus on the influence of the embedding methods on the quality of the final representation. The seven methods we selected implement a broad range of algorithms that are currently used for the analysis of scATAC-seq and scHPTM data. More precisely, ChromSCape_PCA is a simple use of PCA after count per million (CPM) normalization, which serves as baseline. ChromSCape_LSI and Signac implement two variants of the latent semantic indexing (LSI) algorithm, which consists in transforming the count matrix with TF-IDF and applying PCA on that matrix. They have been used to analyze scHPTM data [28, 10], and differ in the fact that ChromSCape_LSI weights the principal components by their eigenvalues, as is standard to do with PCA, while Signac does not and instead whitens the data representation. They implement variants of the algorithm used in Cusanovich2018 [29, 30, 31], which was found with SnapATAC and cisTopic to be among the best methods for scATAC-seq data analysis in [12]. SnapATAC computes the Jaccard similarity between all the cells, and runs kernel PCA on this similarity matrix. cisTopic binarizes the count matrix and then applied latent Dirichlet allocation (LDA) on this modified matrix. Finally SCALE and PeakVI both implement a variational autoencoder (VAE) with a product of Bernoulli likelihood function. They differ in the fact that SCALE uses a mixture of gaussian prior where PeakVI uses a unimodal gaussian prior. Furthermore PeakVI computes corrections for the size factor of each cell as well as for the accessibility of each DNA region. We run all methods with their default parameters (see Methods). In particular, we keep the default number of dimensions for all methods; indeed some methods offer their own heuristics for deciding the number of dimensions, and we did not want to disadvantage them by using a dimension they do not consider optimal. More precisely, PeakVI sets by default the dimension to the square root of the square root of the number of regions, while cisTopic trains model for multiple dimensions and chooses one based on an elbow rule of its evidence lower bound (ELBO). Signac uses a dimension of 50 by default, while SnapATAC, SCALE, and ChromSCape have a default dimension of 10.

Figures 2A and S4 summarize the performance of each embedding method on the different histone PTM marks in the mouse brain and human PBMC datasets, respectively. In those plots, we summarize the performance of each embedding method by reporting the best performance achieved by each embedding method across all possible matrix construction choices, without performing any additional QC processing such as cell or feature selection. This allows us to quantify the best possible result that each embedding method can reach without setting an arbitrary feature engineering pipeline that could advantage some methods over others. We see that the neighbor scores vary roughly in the range 0.05 ∼ 0.35 across methods, datasets and marks. As can be seen in Figures 2B, where we visualize the embeddings obtained by ChromSCape_LSI on different marks on the mouse brain dataset, this corresponds to a fairly good agreement with scRNA-seq embedding in terms of recovering major cell types, particulary for H3K27ac (score=0.302) and H3K4me1 (score=0.321). Interestingly, we observe differences in the neighbor scores of different marks across methods in the mouse brain dataset, with H3K4me1 and H3K27ac (score=0.291*±*0.028 and 0.273 *±* 0.026, respectively) significantly (p=0.008, see Table S4 for all pairwise comparison p-values) higher than H3K9me3 and H3K27me3, and H3K4me3 (score=0.148 *±* 0.040, 0.169 *±* 0.033 and 0.112 *±* 0.035, respectively). Note that this does not necessarily mean that some marks are more informative than others in general, but rather than they are less directly linked to expression than others. A similar trend is visible but weaker on the human PBMC dataset (Figure S4), where in particular the scores on H3K27ac and H3K4me1 are lower than on the mouse brain dataset (scores=0.113 *±* 0.031 and 0.150 ± 0.021, respectively), and only H3K4me1 has a significantly higher scores (p<0.05) than the other marks (see Table S5). This difference between the mouse brain and human PBMC datasets could be caused by the differences in co-assay, by the relative complexity of the cell types, or by the quality of the experiments.

**Figure 2:**
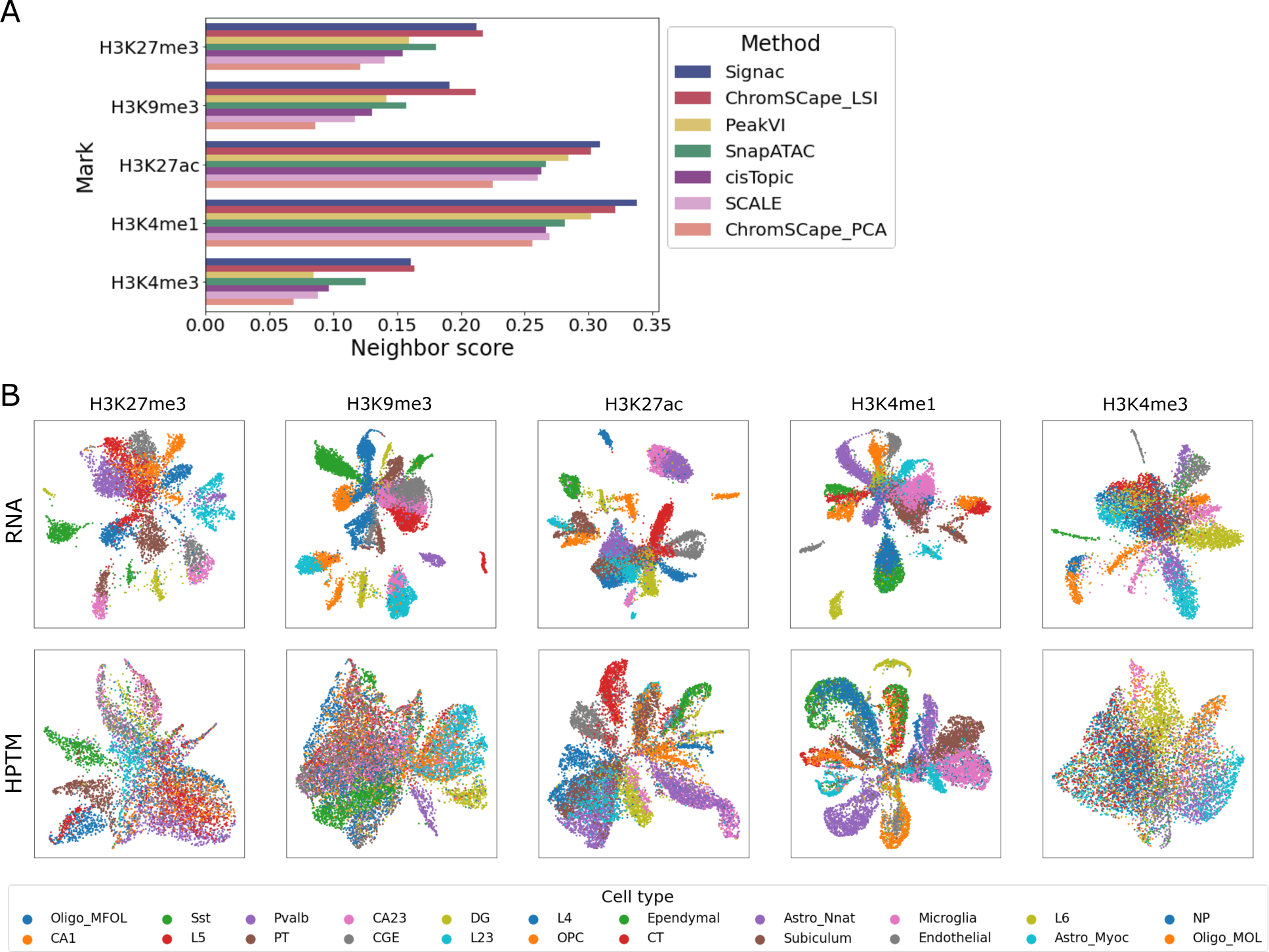
**A**. Best performances of the different representation methods on the mouse brain dataset. **B**. UMAP representation of the different samples in the mouse brain dataset, the first row is the RNA co-assay processed with PCA using the scanpy best practices, the second row is the scHPTM assay processed with ChromSCape_LSI using the matrix construction algorithm with the best neighbor score, both colored by the labels of [9] obtained from the scRNA-seq co-assays.

The performance of each method on each histone PTM mark of the mouse brain datasets is shown in Figure 2 and Table S3. We see that the two best performing methods on the mouse brain datasets are consistently ChromSCape_LSI and Signac, which are significantly better than all other methods (see Table S7 for p-values of pairwise comparisons). They are followed by SnapATAC and PeakVI (except on H3K4me3), then cisTopic, SCALE, and ChromSCape_PCA. SnapATAC is significantly better than cisTopic and SCALE, while ChromSCape_PCA is significantly worse than all other methods. Both top performing methods (ChromSCape_LSI and Signac) implement LSI, suggesting that LSI-based method have an advantage over other approaches. Surprisingly, though, while ChromSCape_LSI also performs well on the human PBMC dataset, Signac does not (Figure S4). This may be due to the lower coverage of the human PBMC dataset than of the mouse brain data, and to the detrimental effect of the whitening operation specific to Signac, as studied in more details in the supplementary text. On the PBMC dataset, ChromSCape_PCA again performs poorly compared to other methods, while the differences between other methods and between marks are overall less pronounced than on the mouse brain dataset.

Since the four methods ChromSCape_PCA, ChromSCape_LSI, Signac and SnapATAC all implement a form of PCA after applying to the count data matrix a specific data transformation, the difference in their performance highlights the importance of this data transformation choice. Simply normalizing the counts by CPM, as ChromSCape_PCA does, leads to poor performances, while normalizing the count data by Jaccard similarity (SnapATAC) or TF-IDF (ChromSCape_LSI and Signac) is consistently better. This seems to be specific to scHTPM, since methods using CPM normalization are competitive with the ones using TF-IDF or kernel PCA on the Jaccard similarity on scATAC-seq data [12].

We also find that cisTopic is not among the best performing methods for the analysis of scHPTM, while it was identified by [12] as one of the best tools for analysing scATAC-seq. On the other hand, LSI is extremely competitive for both modalities. This shows that while scHPTM and scATAC-seq have some similarities, one should be careful before extrapolating good practices from one modality to the other. Finally, the more recent VAE-based methods, PeakVI and SCALE, are overall not competitive with the more classical LSI-based ones. As we show below, this may be due to the relatively small size of the datasets used.

### The count matrix construction strongly influences the quality of the representation

We now investigate the influence of the count matrix construction method (i.e., how the raw reads are mapped to regions) to obtain relevant embeddings of scHTPM datasets. For that purpose, we explore the performance of the different embedding methods as a function of the matrix construction parameter, again without further preprocessing such as cell or feature selection. We show the results in Figures 3 and S5 for the mouse brain and human PBMC datasets, respectively.

**Figure 3:**
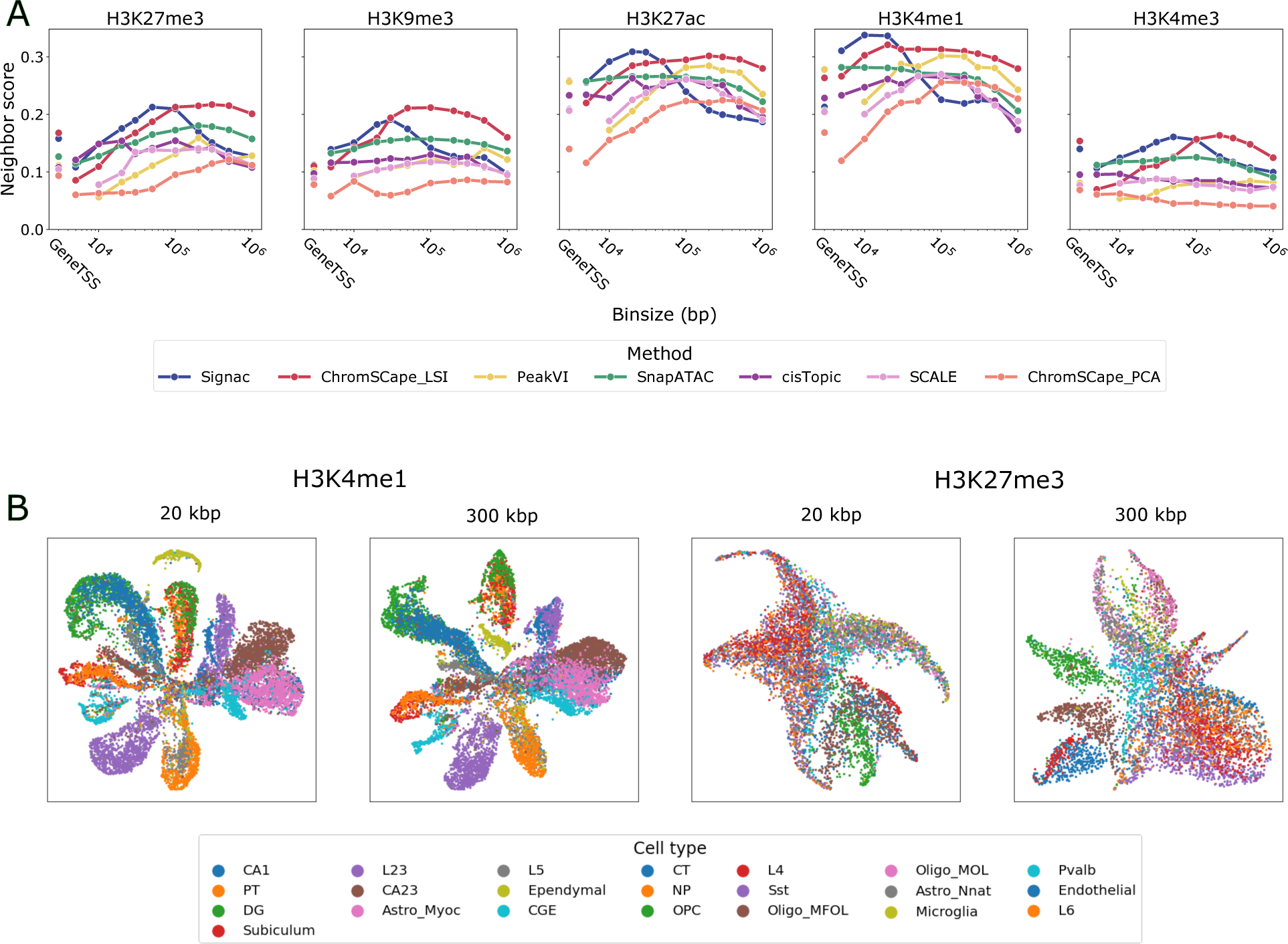
**A**. Performances of the 7 dimension reduction algorithms on the 5 marks in the mouse brain dataset, as a function of the matrix construction. **B**. UMAP projecttion of H3K4me1 and H3K27me3 using ChromsSCape_LSI using bins of 20kbp and 300kbp, colored by the labels of [9] obtained from the scRNA-seq co-assays.

We see that matrix construction has overall a strong influence on the quality of the representations. For most methods and marks, the performance first increases when the bin size increases, then decrease after a peak. This effect is more pronounced on the mouse brain data, and in particular for repressive marks (H3K27me3 and H3K9me3). In order to quantify this effect, we report the ratios between the best and worst performing matrix construction for each method and mark in Table S10 for the mouse brain dataset and in Table S11 for the human PBMC dataset. In the human PBMC dataset, we can see that the ratio between the best and worst feature engineering can reach up to 7.64 (PeakVI on H3K4me1), this is mostly due to the very poor performances of using a GeneTSS annotation on this dataset as can be seen in Fig S5.

In the mouse brain dataset, we can see that this ratio is on average above 2 for H3K27me3 in Fig S9 and reaches 2.8 in the case of PeakVI. The lowest ratio is 1.2 (ChromSCape_LSI on H3K4me1), which is still an increase in performance of 20%. While that ratio is on average higher for the best performing methods (ChromSCape_LSI and Signac), it is mostly due to the fact that their best performances are higher than the other methods, more than it is due to an extreme sensitivity to matrix construction. Indeed we can see that for all marks, ChromSCape_LSI has a very large range of matrix construction parameters that are extremely competitive. We can also note that by choosing an average performing method (e.g. SnapATAC or PeakVI) and an appropriate matrix construction parameter, we can always beat the best performing methods (ChromSCape_LSI or Signac) if they are run with a suboptimal parameter for matrix construction.

We see on the mouse brain dataset that performances reach a level close to their maximum for smaller bin sizes for enhancing marks (H3K27ac, H3K4me1 and H3K4me3) than for repressive marks (H3K27me3 and H3K9me3), and that, except for Signac, the range of appropriate bin size is relatively large (e.g. 50kbp-1000kbp for H3K27me3 or 10kbp-200kbp for H3K4me1). Furthermore, except for Signac, that range is relatively stable across methods for each bin size. We investigate in more details the reason why Signac behaves so distinctively in the supplementary text (Fig S1), and show in particular that the fact that it uses a whitening step and a relatively high embedding dimension by default makes it might capture more noise for larger bin sizes.

We observe that using the GeneTSS annotation is usually not competitive compared to using an appropriate bin size. The fact that H3K4me3 is an exception to that rule is consistent with the fact that this mark is known to be particularly enriched around genes and TSSs. We can also see in Fig S5 that the PseudoBulk annotation is also generally not competitive, with a less pronounced effect for H3K4me1 and H3K4me3. This is consistent with the fact that these marks tend to have small peaks, which are easier to identify with peak calling algorithms than larger ones.

It is interesting to note that the range of appropriate bin sizes for optimal representations usually includes 100kbp and can even go up to 500kbp, which would a priori be considered too large to keep biological relevant information. In particular in [8], the authors made the choice of 5kbp for H3K4me3 and 50kbp for H3K27me3, while in [9] the authors chose 5kbp for all marks, except for H3k4me3 for which it was 1kbp. Here we find that, to reach a maximal concordance between epigenomic and transcriptomic embeddings, bin sizes one or two orders of magnitude larger than the ones used in previous studies are still competitive. This is likely due in part to the fact that the coverage per cell is so low that taking smaller bins introduces too much noise in the matrix, and to the fact that genes are not randomly distributed in the genome, and tend to cluster into groups of co-expressed genes [32, 33],. We can also observe from Fig 3 that LSI based methods such as Seurat can achieve good performances in the lower bin sizes regime (as well as ChromSCape_LSI as shown in the supplementary text).

### Selecting high coverage cells has a modest positive impact on the representation

In a standard QC pipeline, poorly covered cells can be filtered out before performing dimensionality reduction and subsequent analysis on the highest quality cells. Such selection step often leads to a trade-off between keeping a high number of cells to maximise the discovery rate of rare cell states, and keeping only highly-covered cells to maximise the quality of the embedding. We now assess how selection of cells based on coverage affects the quality of the embedding, by applying different thresholds for coverage selection and measuring neighbor scores across methods.

As shown on Fig 4, there is overall a modest gain in performance when applying more stringent QC criteria on cell coverage. Across histone marks, we observe a maximum gain of 15% and 13% in performance for H3K4me1 when using the best performing methods ChromSCape_LSI or Signac respectively (Table S12). Across methods, we observe that the highest gains in performances are observed for the low performing methods identified above Table S13. ChromSCape_PCA and SCALE benefit from a 41% and 21% gain respectively whereas ChromSCape_LSI only benefits from an average 8% gain. In summary, filtering out cells with low coverage has little impact on the quality of the representation, while reducing the probability to capture rare cells.

**Figure 4:**
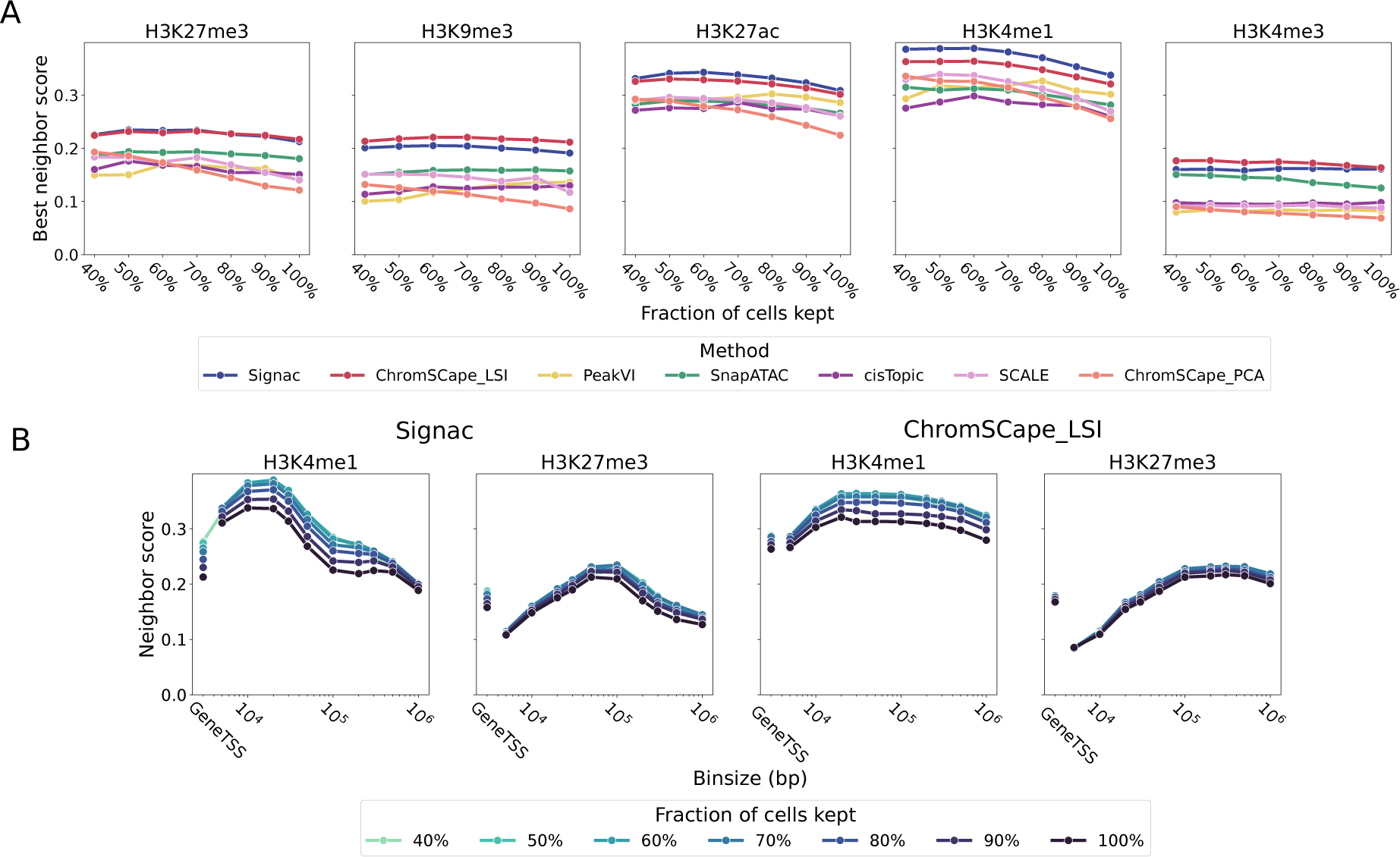
**A**. Each point corresponds to the best performance across matrix construction of a given method and a given coverage threshold, for the 7 methods, 5 marks, and 7 coverage conditions. **B**. Performances of Signac and ChromSCape_LSI as a function of matrix construction on H3K4me1 and H3K27me3 for different coverage thresholds.

A related question important to prepare the experiments is, independently of the number of cells filtered out, to clarify the impact of average cell coverage. As studied in Supplementary text, we observe that for a given number of cells, the average coverage per cell is strongly positively correlated with the quality of the representation for most marks and methods. This confirms that low cell coverage is currently one of the main reasons behind the difficulty to analyze scHTPM data and to capture a robust biological representation of each cell.

### Feature selection decreases the quality of the embedding

Another QC criteria used in single-cell analysis is the selection of features - genomic regions for single-cell epigenomics datasets - prior to dimensionality reduction. Two standard approaches are (i) the selection of regions with the highest coverage or (ii) the selection of regions that have a highly variable enrichment score across cells. Such a selection step is relatively common, but there is currently no consensus for scHPTM analysis on whether such selection is beneficial, and which of the two methods is optimal.

To address this question, we compare the maximal neighborhood scores for all methods with various feature selection thresholds, when we select features based on variability (HVG) or coverage. The results are shown on Figures 5.A and S6 respectively, for the mouse brain dataset. We observe consistently that feature selection is generally detrimental to the performances, in the sense that for both methods, the more regions we keep the better the performances are. As shown on Figure 5.B for Signac and ChromSCape_LSI, this trend is in fact not only true when we look at the best performance reached over different bin sizes in the matrix construction step, but also when we look at each bin size individually.

**Figure 5:**
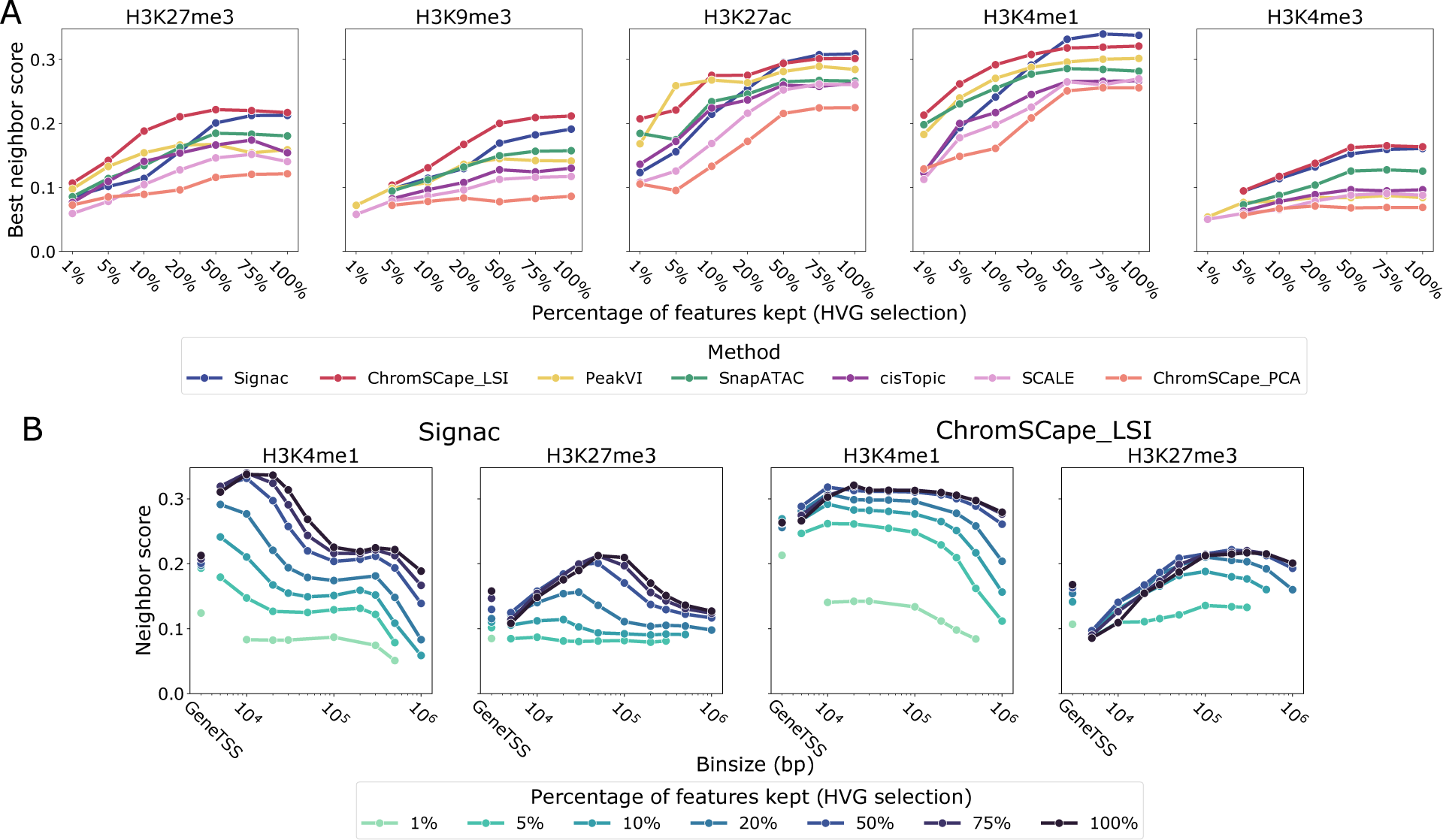
Role of feature selection, using the Highly Variable Gene (HVG) method used for scRNA-seq on the mouse brain dataset.**A**. Each point corresponds to the best performance across matrix construction of a given method and a given percentage of features kept, for the 7 methods, 5 marks, and 7 feature selection conditions. **B**. Performances of Signac and ChromSCape_LSI as a function of matrix construction on H3K4me1 and H3K27me3 for different feature selection thresholds.

Feature selection has been shown to increase performances for scRNA-seq in [13] and is part of the guidelines for scATAC-seq [24]. Our results show that, contrary to scRNA-seq and scATAC-seq, feature selection is detrimental to the analysis of scHPTM data, and we therefore recommend not to use it.

### Performances reach a plateau near 6000 cells

While computational parameters can have an important role in the quality of the representation [13], experimental ones also have a strong influence. In this section we look at the role of the number of cells on such representations, in order to help practitioners design their experiments. For that purpose, we systematically downsample each dataset by randomly selecting a subset of cells of various size, and assess the quality of the representation obtained from the downsampled datasets. We show on Figure 6.A the best performance reached across matrix construction for each method on each mark, as a function of the size of the downsampled dataset, for the mouse brain dataset. We further add a finer grained sweep over dataset size for ChromSCape_LSI, by increasing the size of the datset by 500 cells per step as can be seen in Fig 6.B.

**Figure 6:**
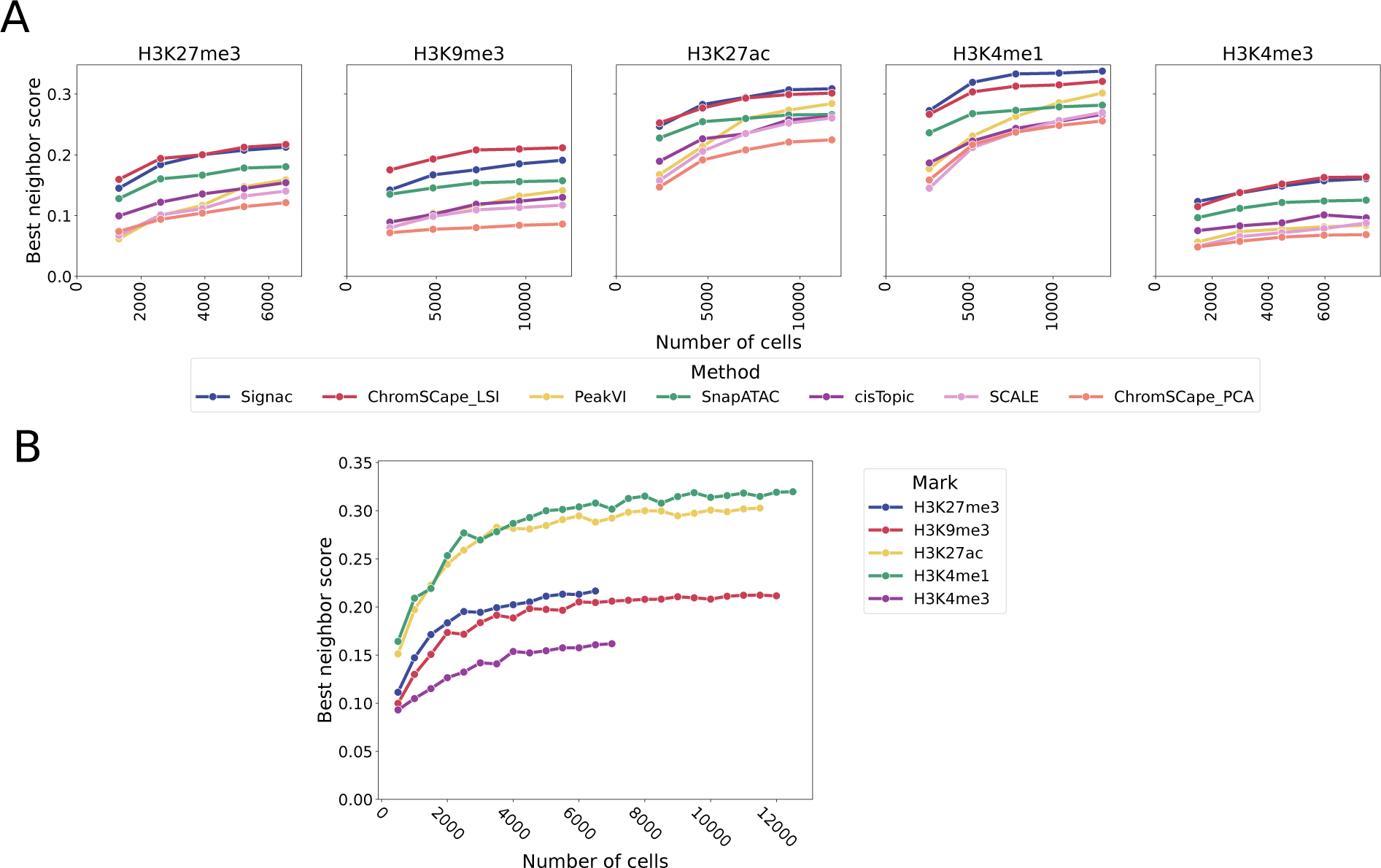
Effect of downsampling uniformly at random the number of cells in the experiment. Each point corresponds to the best performance across matrix construction. **A**. Performances the 7 methods, on the 5 marks of the mouse brain dataset and on 5 sizes of dataset (by increase of 20% of the dataset size). **B**. Performances of ChromSCape_LSI on the 5 marks, using an increase of 500 cells per step.

We see that all methods, on all datasets, benefit from an increase in the number of cells. However, it is interesting to note that the benefits resulting from a larger number of cells diminish as the number of cells increases. Indeed we can observe that the performances increase quickly up to ∼6,000 cells, and then only keep increasing at a much smaller rate. PeakVI is an exception to that observation, and we can see that its performances have not yet reached this plateau, see Table S15. This is consistent with the intuition that deep learning based models require a large amount of data to achieve their full performances, and in the datasets used in this paper, this full performance does not seem to have been achieved. The gains in performances are also quite important, with an average increase of 34% by increasing the number of cells by 150%, and 18% by increasing the number of cells by 66%.

On the other hand the more standard methods, such as LSI or kernel PCA, reach their peak performances around 6,000 cells, and only gain an average of 5% in performances by going from 6,000 to 10,000 cells. Since these methods are the best performing ones in regime tested in this paper (less than 12,000 cells), it means that practitioners can sequence less cells while keeping relatively good performances. The case of ChromSCape_LSI is shown in more details in Fig 6.B, where the dimishing return effect of adding more cells is very pronounced. It also allows us to confirm that the difference in performances between the enhancing and repressive marks is not due to the number of cells present in the datasets, as we see a clear separation of 3 groups: H3K27ac and H3K4me1 having the best performances, H3K9me3 and H3K27me3 following them, and finally H3K4me3 having the worst performances. The plateauing effect can also be seen for all of these marks, leading us to believe that it is not specific to just some marks.

It is however very possible that given more cells (>12,000), PeakVI and SCALE could outperform these methods and lead to better representations. This would be consistent with the behaviour of deep learning-based methods on other modalities such as text of images, even though we can only conjecture that this would happen.

While the increase in the number of cells leads to observable and consistent gains in the quality of the representation, it is noteworthy that these gains have a lower influence than the use of an optimal matrix construction algorithm. It is also important to note that the performances of the current best methods do not strongly benefit from such an increase in the number of cells as can be seen in Table S14, meaning that practitioners may work on relatively small samples while maintaining state-of-the-art performances.

## Discussion

In this paper we studied the role of experimental parameters, cell and feature selection, matrix construction, and dimension reduction on the quality of the representation from scHPTM datasets. We decided to focus on the quality of the dimension reduction as it is generally the input of most downstream tasks such as clustering, cell type identification, differential enrichment or trajectory inference. A good representation is thus beneficial to all these tasks.

Unlike other benchmarks [12, 13, 15] we decided not to measure the quality of the representation based on the ability of clustering algorithms to retrieve known cell types. This is due to the lack of high quality labels for scHPTM data. One possibility for obtaining labels could be to use the labels derived from the co-assay, however this would rely on computational methods instead of an orthogonal protocol and thus would not allow us to truly measure the quality of the representation. An alternative would be to use FACS-sorted data, whose label are not computational in nature, however no such data exists that presents interesting complexity to our knowledge. FACS-sorted label also suffer from being discrete in nature, which may not be informative to how well we could represent continuous state transition in cell differentiation. The last alternative would be to use simulation data, but not only is there no such simulation tool accepted in the community, but also performances on simulated data may not be transferable to real data. Instead we decided to evaluate how well the representation in scHPTM locally agrees with a reference co-assay. This approach allows us to be independent of labels, as well as working with potentially continuous cell states. Yet when using this score, we make the assumption that cells that have similar epigenomic landscapes, measured by scHPTM, should at least locally display similar RNA or protein expression patterns. While we know that this assumption holds well for enhancer marks (e.g. H3K4me1 or H3K27ac), it might suffer some exceptions for repressive marks (H3K27me3, H3K9me3). Overall, looking at the available datasets and prior biological knowledge, it appears reasonable and the best we can have without proper labels. We can also note that this approach has already been successfully used in evaluating scATAC-seq pipelines in [19, 18].

While we expected the choice of matrix construction algorithm to have an impact, that impact is larger than what we expected a priori. Indeed as is shown in Table S8 the performances using the best bin size can be up to 80% better than using the worst one. Surprisingly the ranges of bin size are larger than what could be expected a priori, and we were also surprised to find that enhancing marks such as H3K4me1 benefited from very large bin sizes (up to 200kbp) despite being known to have small peaks (in the order of a few kbps [17]). Yet, at a bin size of more than 50kbp, embeddings will not rely on local epigenomic enrichments, such as the ones observed for enhancers. The coverage of current scHPTM datasets might not be sufficient to produce reliable embeddings from small bins, and may thereby be unable to distinguish cell states distinguished by a few local differences. Identifying differences in enrichment for smaller regions than the bin size used for the embeddings can however be done by using a bin size more appropriate for the biology of the studied mark when running differential enrichment analysis, while using the clusters obtained from the embedding. We can also consider the fact that our evaluation relies on the similarities between gene expression and gene regulation, which may be biased by the known existence of co-expressed gene clusters throughout the genome [32, 33]. With bin sizes over 100kbp, we might be robustly detecting such co-regulated gene clusters. The fact that GeneTSS and PseudoBulk annotations were in general not competitive is also not something that was not previously rigorously established in the literature to the best of our knowledge.

It was also interesting to note that, except for PeakVI, the performances of the different methods tend to stagnate when increasing the number of cells. This is likely due to the relatively low complexity of the models used. More complex models such as PeakVI or SCALE did not manage to outperform these low complexity ones in our experiments. One could imagine that these models could show better performances with larger datasets, such as cell atlases, but they do not seem appropriate for experiments as they are currently designed.

On the other hand, we found that the performances of all methods largely benefited from being run on high coverage cells, and that these performances did not stagnate on the available data, suggesting that future improvements of protocols - increasing coverage, such as in [34] - will surely provide additional information and granularity to refine embeddings.

We were also surprised to observe that feature selection, using either a variance or a coverage criteria, almost always had a negative impact on the performances. This may be due to the excessively low coverage per cells compared to other protocols where this procedure can be beneficial (see [13] for scRNA-seq).

To the best of our knowledge, this manuscript provides the first comprehensive study on how to both design the experiment, build the matrix, and analyse scHPTM data. We hope that the large effect of matrix construction that we were able to identify will lead the community to pay more attention to this crucial, if overlooked step. Furthermore by testing the algorithms and pipelines most likely to work on scHPTM data, we hope to save the community some time by avoiding having to discover which already existing algorithms work best.

## Material and Methods

### Matrix construction

We downloaded the mouse brain dataset from GSE152020 (https://www.ncbi.nlm.nih.gov/geo/query/acc.cgi?acc=GSE152020). The data come in count matrix format, with 5kbp bins for all marks except for H3K4me3 which is in 1kbp bins. The larger bin sizes were obtained by merging the original bins together to form new bins using a custom script, available at https://github.com/vallotlab/benchmark_scepigenomics. The GeneTSS annotation comes from the ChromSCape package, and the matrix was done by merging the bins containing the regions in that annotation using the same custom script. We keep all the cells present in that matrix, as the original authors already applied QC steps on the cells.

The human PBMC dataset data was downloaded from https://zenodo.org/record/5504061, the data was processed from the fragment files. We used ChromSCape for generating 5kbp matrices and then used our custom script to generate the other matrices similarly to the mouse brain dataset. The PseudoBulk annotation was obtained by turning the fragment files into bams, calling the peaks using MACS2, and then merging them using bedtools. We then select only the cells used in the original paper analysis, by keeping only the barcodes present in the rds objects on zenodo.

### In silico modifications

Using the matrices generated in the previous section, we modified them in order to both simulate experimental conditions, as well as apply standard bioinformatics steps. Feature selection was done using Seurat’s `FindVariableFeatures` for the HVG selection and ChromSCape’s `find_top_features` for the top regions selection, ran using our `filter_sce.R` script. For selecting only the high coverage cells we sorted cells by coverage, and kept only the most covered ones, the relevant script is `filter_cells_quality.R`. For studying the role of the number of cells, we sampled cells at ramdom without replacement from the matrice, the relevant script is `sample_cells.R`.

### scRNA-seq and CITE-seq processing

The scRNA-seq matrix for the mouse brain dataset was processed using the scanpy [21] package, and following their best practice notebook (https://scanpy-tutorials.readthedocs.io/en/latest/pbmc3k.html). We have previously shown in [13] that the algorithms used in that package are robust and perform well. The CITE-seq matrix for the human PBMC dataset was extracted from the rds objects and processed with standard PCA.

### Representations for scHPTM

For computing the representations using the different methods, we used the implementation from the original packages, except for SnapATAC for which we used the reimplementation of [12] as it allowed a nicer API for running a large number of jobs, their implementation can be found on their github https://github.com/pinellolab/scATAC-benchmarking. For cisTopic, we ran the `runWarpLDAModels` method from the cisTopic Bioconductor package (version 0.3.0) and followed the steps from [12]. For Signac we followed the scATAC-seq best practices vignette (https://satijalab.org/signac/articles/pbmc_vignette.html) and used the Signac CRAN package (version 1.7.0). For ChromSCape_LSI and ChromSCape_PCA we processed the matrix with the `tpm_norm` and `TFIDF` methods respectively, then called the `pca` method, and removed the first principal component, all the methods were callled from the ChromSCape Bioconductor package (version 1.6.0). For PeakVI we followed the tutorial on the package website https://docs.scvi-tools.org/en/0.15.1/tutorials/notebooks/PeakVI.html using the scvi-tools (version 0.15.1) [35] pip package. Since SCALE did not have an API for calling their model, we modified the `main.py` script from the `scale` python package (version 1.1.0), so that it does not remove cells.

The scripts for processing used for all R methods ate in the `R_analysis.R` script, PeakVI and SCALE are respectively `peakVI_process.py` and `scale_process.py` scripts.

The R methods were run on CPUs with 16 cores and 32GB of RAM, the deep learning ones (PeakVI and SCALE) on V100 GPUs with an additional 2 cores CPU.

### Neighbor score computation

To compute the neighbor score of an scHPTM representation, we first compute the *k* nearest neighbor graphs (kNNG) for values of *k* ranging from 0.1% up to 10% of the cells present in the dataset. We then compute the representation for the second modality using scanpy [21], whose algorithm (PCA) has been identified in [13] as being the most reliable for achieving good representations for scRNA-seq. We then compute kNNG on this second representation, count the number of common neighbors in the kNNG for each cell, divide by *k*, and average over the cells. This gives a score between 0 and 1, where 1 means that the two representations perfectly agree on which cells are similar, and a score of 0 means complete disagreement. We further average that score across the various values of *k*, which were selected to be 0.1%, 0.3%, 0.5%, 1%, 3%, 5% and 10% of the cells contained in the assay, in order to take into account the multiple possible levels of similarity. Two completely random representations would have a score of 0.05 given the values of *k* that we selected.

### Compute time

The average time for computing an embedding per method is presented in Table 2. The scores computation took an additional 30 minutes per seven embeddings. This resulted in about 1 year equivalent of CPU/GPU time to run the full benchmark.

**Table 2:** Average time for computing an embedding per method, small matrix is large bin size (>200kbp), small matrix is <100kbp.

|  | architecture | small matrix | large matrix |
| --- | --- | --- | --- |
| Signac | CPU 16 cores | 3 minutes | 6 minutes |
| ChromScape_LSI | CPU 16 cores | 3 minutes | 6 minutes |
| ChromScape_PCA | CPU 16 cores | 3 minutes | 6 minutes |
| SnapATAC | CPU 16 cores | 10 minutes | 40 minutes |
| cisTopic | CPU 16 cores | 15 minutes | 60 minutes |
| PeakVI | GPU V100 | 10 minutes (early stopping) | 3 hours |
| SCALE | GPU V100 | 10 minutes (early stopping) | 3 hours |

## Conflict of interest

The authors declare that they have no competing interests.

## Supplementary material

### Supplementary text

#### Role of whitening in LSI

**Figure S1:**
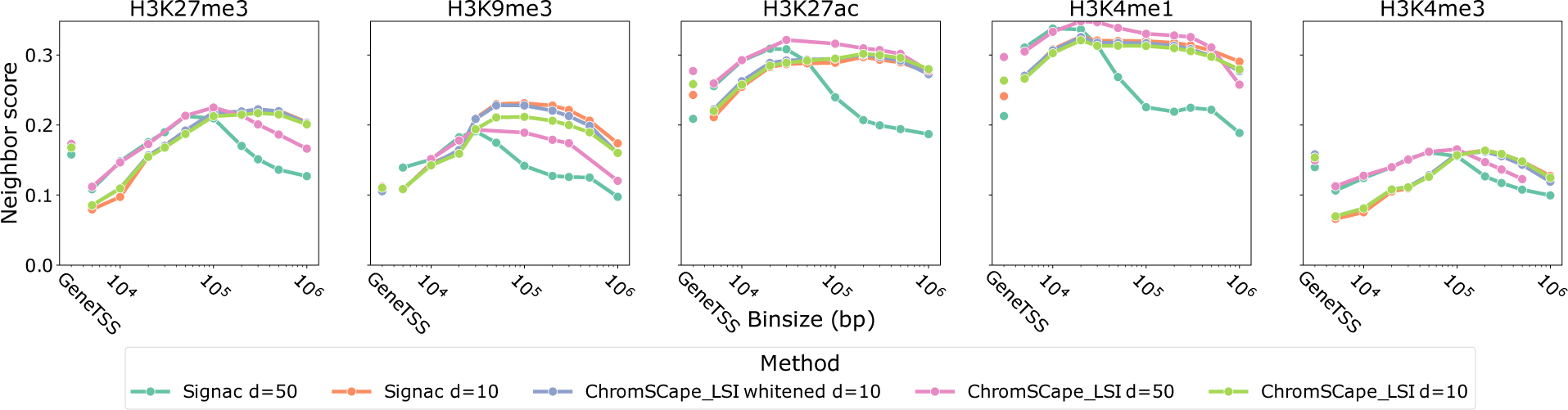
Performances of Signac, ChromSCape_LSI, and modified ChromSCape_LSI on the 5 marks in the mouse brain dataset, as a function of the matrix construction

**Figure S2:**
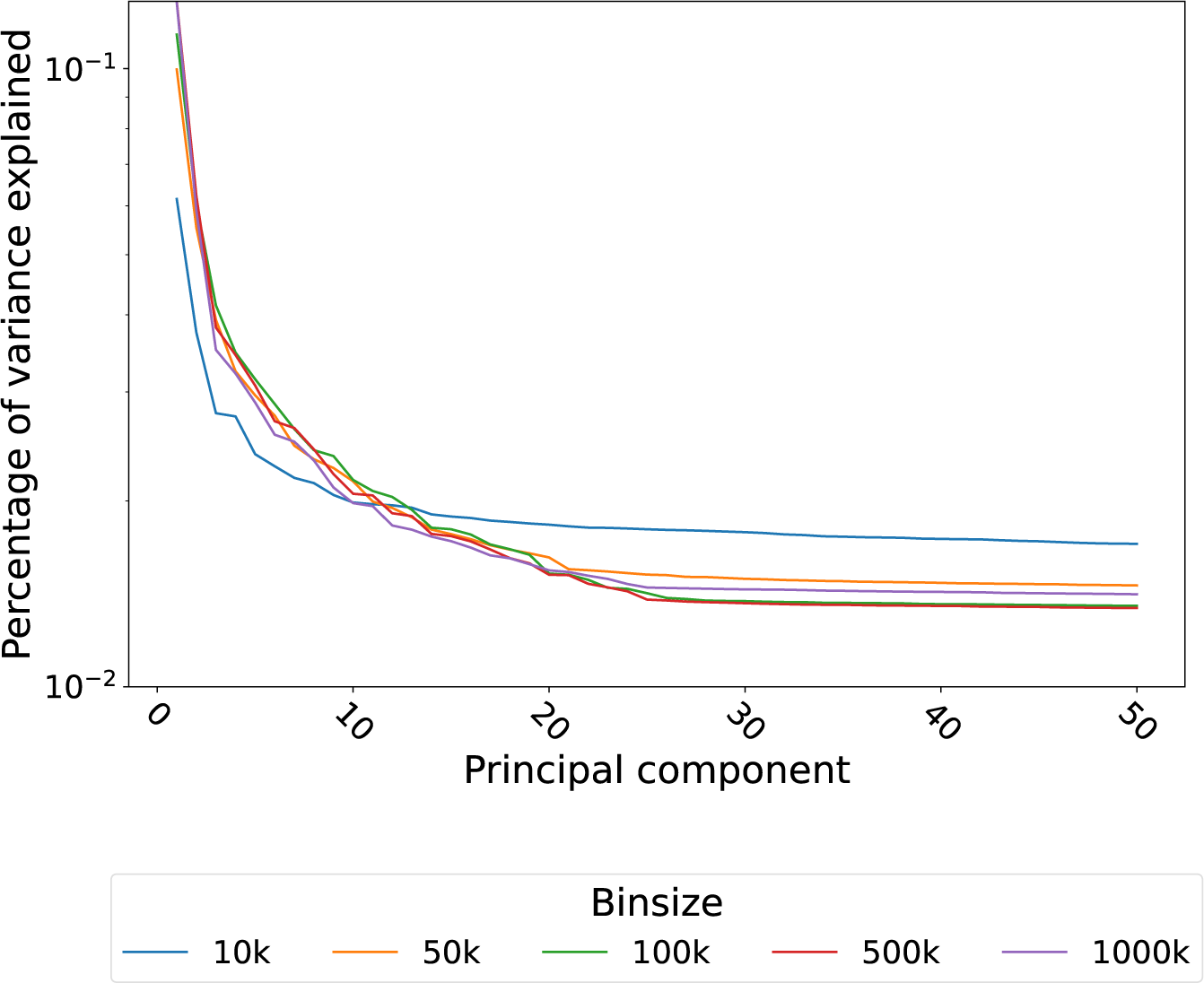
Percentage of variance explained by each principal component of LSI on H3K4me1 of the mouse brain dataset.

We could see in Fig 3 that Signac’s best performances are achieved for a smaller bin size than the other methods, it is especially surprising because it is supposed to implement the same algorithm as ChromSCape_LSI. In this section we investigate the reason behind this difference.

A close look at the implementation of Signac shows that it does not implement the standard LSI algorithm, but instead a whitened version of it; the principal components (PC) are not weighted by their explained variance. Furthermore the default dimension used is 50 instead of 10 for ChromSCape_LSI. Both methods also remove the first PC, as it is assumed to be mostly driven by the coverage per cell. In order to understand the cause of the shift in optimal bin size we ran Signac with dimension 10, and ran ChromSCape_LSI at dimension 50 and modified it to use whitening. By comparing the three conditions with a dimension of 10, we can see that the difference in performances between Signac and ChromSCape_LSI in the previous sections can mostly be explained by the number of PCs, indeed except in the case of H3K9me3 the performances of the two are almost identical as can be seen in Fig S1.

However we can see that in higher dimensions, 50 PCs, the performances are very different. While the top performances are comparable, with a slight advantage for ChromSCape_LSI, Signac has a much tighter range of bin sizes with good performances. This can be explained by the fact that in large dimensions, the later PCs explain less variance than the early ones, and not weighing appropriately induces noise. We can see that when the bin size is small, the explained variance per PC is less concentrated in the first PCs than in the later ones, this is shown in Fig S2. This explains why Signac has a tighter range of good bin sizes, as its whitening makes the later PCs as important as the first ones.

We thus recommend not whitening when using LSI, as it makes the method less robust to the choice of bin size.

### Trade-off between coverage and cell number

**Figure S3:**
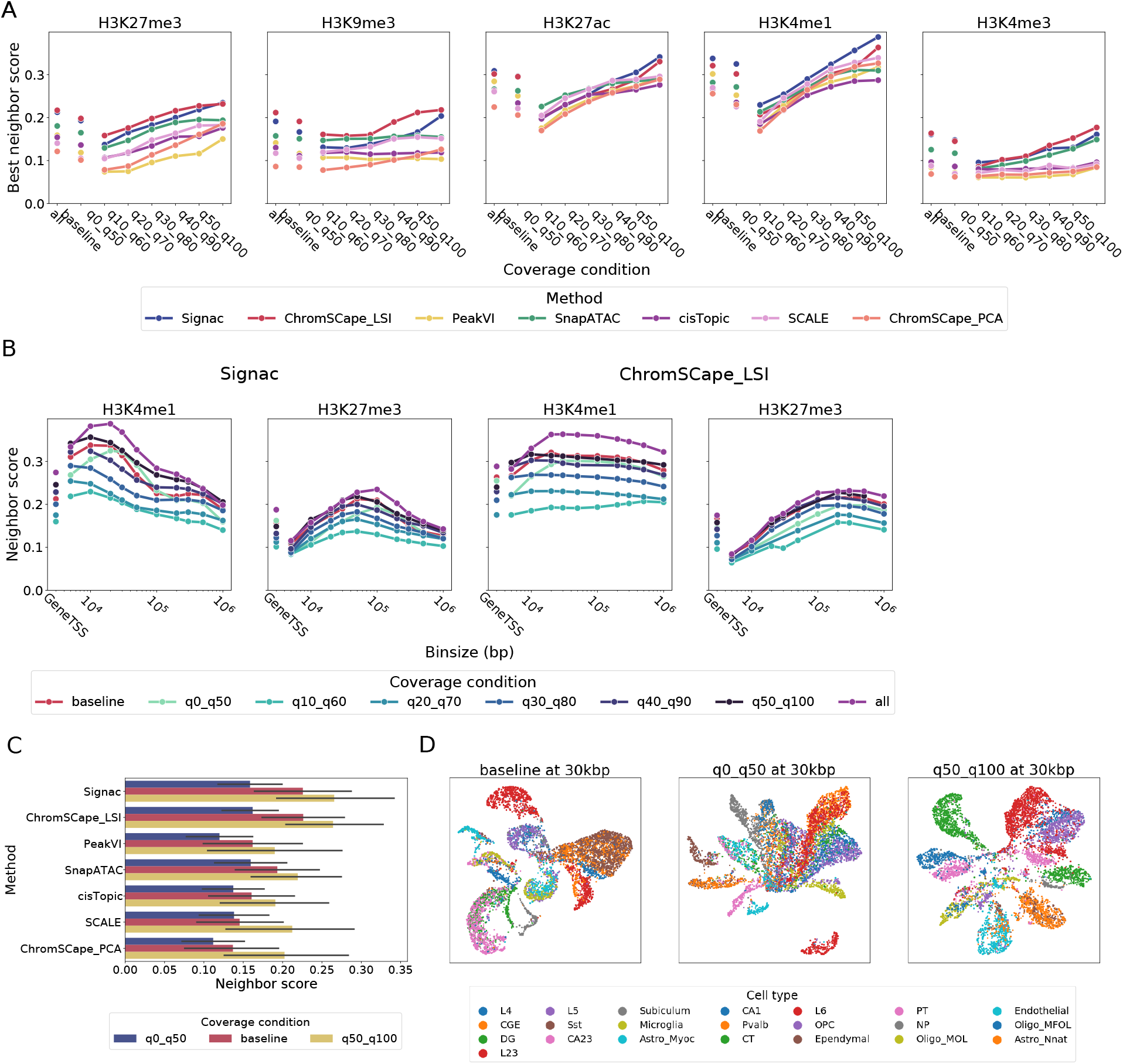
Study of the effect of cell coverage on the performances of the representations. The all condition contains all the cell as a reference, the baseline condition contains only 50% of the cells uniformly sampled at random, the other 6 conditions contain 50% of the cells, but are sorted by coverage. We order the cells by how much reads they contain and take all the cells from the bottom n% up to *n* + 50% in order to have the same amount of cells in all conditions. **A**Best performance across matrix construction, measured for all of the 7 methods, 5 marks of the mouse brain dataset and 8 coverage conditions. **B**. Performances of Signac and ChromSCape_LSI on H3K4me1 and H3K27me3 as a function of matrix construction. **C**. Average best performance of the 7 methods across all marks, for the lowest covered cells, random cells, and highest covered cells. **D**. UMAP projection of H3K4me1 at different coverage qualities, using ChromSCape_LSI across at 30kbp, colored by the labels of [9] obtained from the scRNA-seq co-assays.

While experimentalists do not control the depth of sequencing per cell with the current technologies, we felt that studying how increasing the coverage would impact the quality of the representations to be of interest. Indeed we currently do not know whether there would be a benefit from such an increase, and if there is one, how strong its effect would be.

In order to evaluate the effect of coverage, we select 50% of the cells, but constrain them to have similar coverage. For example in the q0_q50 condition we take the 50% cells with the lowest coverage per cell, and in q50_q100 we take the cells with the highest coverage. In a more general way, we sort the cells by coverage per cell, and select the cells whose coverage falls between the *n*-th percentile, and *n* + 50-th percentile. This allows us to have all conditions with the same amount of cells, and just study the effect of coverage. We also have a condition where we just sample half of the cells at random, including all coverage, in order to have a baseline to compare against. That protocol is summarized in Fig S3. This approach of sampling the cells by coverage instead of downsampling the reads per cell has the advantage that it does not make any assumption on the data generation process. Indeed here all the observed cells are real cells, instead of cells that are modified with computational means.

First we can observe that the performances of all methods increase as we increase the coverage, which was expected. However unlike the number of cells in an experiment seen in Fig 6, the positive effect of more reads per cells does not plateau and is almost a straight line in the case of H3K4me1 as we can see in Fig S3. If we look at the differences in performances between the least and most covered cells, summarized by mark in Table S16 and by method in Table S17, we can see an increase of at least 35%. That increase even goes as far as 107% for H3K4me3 with ChromSCape_LSI. Looking at the difference between the baseline and the high coverage cells, that increase is still in the order of 15%. It is interesting to notice that this effect on performances is larger than the one obtained by an increase in the number of cells, furthermore as we can see in Fig S3 this gain is not yet completely achieved in our protocol. That effect is specifically noticeable for H3K27me3. This also agrees with the results on the section studying the role of selecting cells by coverage, where we identified that the marks benefiting the most from this selection were H3K27me3, H3K27ac, and H3K4me1.

The effect of coverage is also consistent across methods, with Signac, ChromSCape_LSI an ChromSCape_PCA benefiting the most form this increase in coverage, above 60%. The first two already being the best performing methods allows to fully reap the benefits from a high coverage.

## Supplementary tables

**Table S1:** Percentage of successful runs on the mouse brain data.

|  | H3K27me3 | H3K9me3 | H3K27ac | H3K4me1 | H3K4me3 |
| --- | --- | --- | --- | --- | --- |
| Signac | 88.2% | 88.5% | 90.3% | 93.6% | 84.8% |
| ChromSCape_LSI | 80.0% | 80.3% | 78.8% | 93.9% | 78.2% |
| PeakVI | 86.1% | 87.9% | 85.2% | 84.5% | 77.3% |
| SnapATAC | 88.2% | 89.1% | 90.6% | 94.8% | 84.8% |
| cisTopic | 88.2% | 88.5% | 90.3% | 93.6% | 84.8% |
| SCALE | 86.1% | 87.3% | 90.6% | 92.1% | 85.2% |
| ChromSCape_PCA | 88.2% | 88.5% | 90.3% | 93.6% | 84.8% |

**Table S2:**
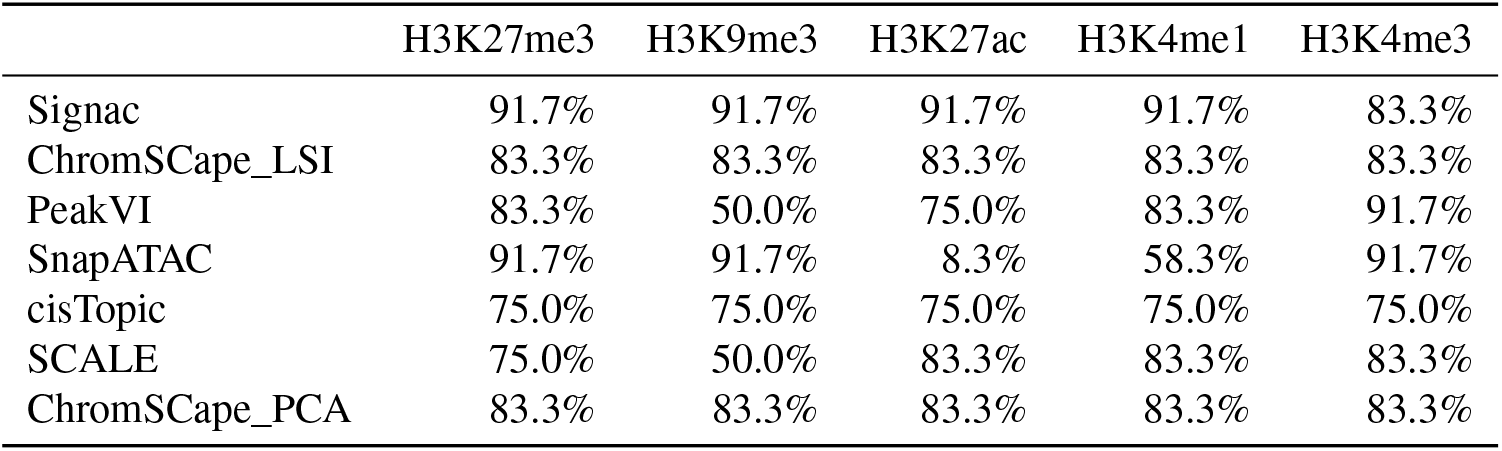
Percentage of successful runs on the human PBMC data.

|  | H3K27me3 | H3K9me3 | H3K27ac | H3K4me1 | H3K4me3 |
| --- | --- | --- | --- | --- | --- |
| Signac | 91.7% | 91.7% | 91.7% | 91.7% | 83.3% |
| ChromSCape_LSI | 83.3% | 83.3% | 83.3% | 83.3% | 83.3% |
| PeakVI | 83.3% | 50.0% | 75.0% | 83.3% | 91.7% |
| SnapATAC | 91.7% | 91.7% | 8.3% | 58.3% | 91.7% |
| cisTopic | 75.0% | 75.0% | 75.0% | 75.0% | 75.0% |
| SCALE | 75.0% | 50.0% | 83.3% | 83.3% | 83.3% |
| ChromSCape_PCA | 83.3% | 83.3% | 83.3% | 83.3% | 83.3% |

**Table S3:**
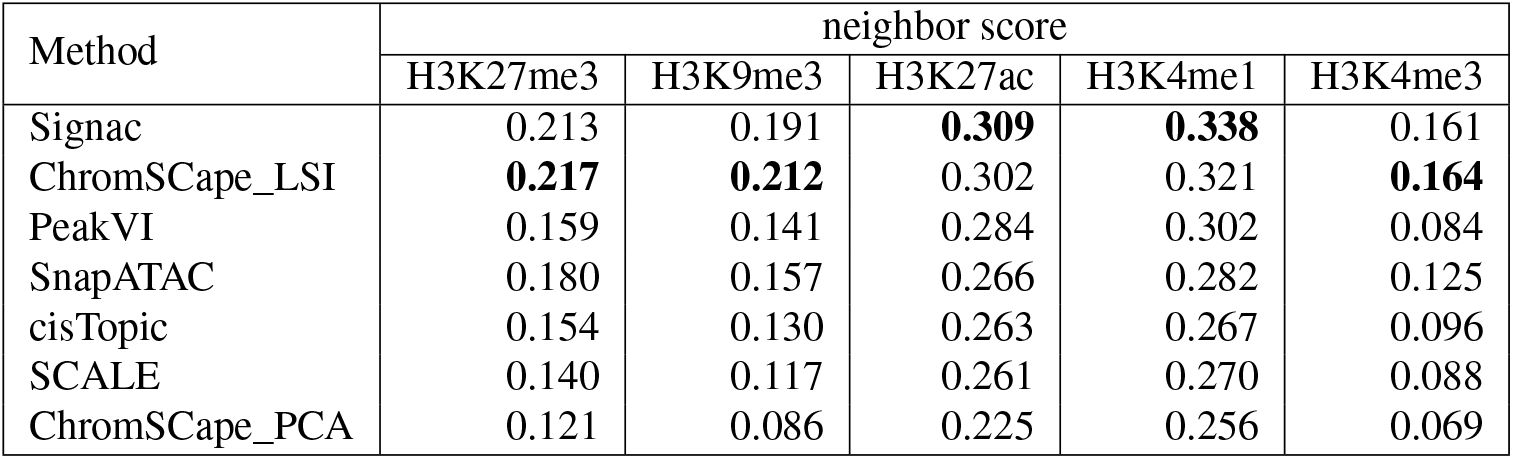
Best performance of each method across feature engineering methods on the mouse brain dataset, the best performing method for each mark is bolded.

| Method | neighbor score |  |  |  |  |
| --- | --- | --- | --- | --- | --- |
|  | H3K27me3 | H3K9me3 | H3K27ac | H3K4me1 | H3K4me3 |
| Signac | 0.213 | 0.191 | **0.309** | **0.338** | 0.161 |
| ChromSCape_LSI | **0.217** | **0.212** | 0.302 | 0.321 | **0.164** |
| PeakVI | 0.159 | 0.141 | 0.284 | 0.302 | 0.084 |
| SnapATAC | 0.180 | 0.157 | 0.266 | 0.282 | 0.125 |
| cisTopic | 0.154 | 0.130 | 0.263 | 0.267 | 0.096 |
| SCALE | 0.140 | 0.117 | 0.261 | 0.270 | 0.088 |
| ChromSCape_PCA | 0.121 | 0.086 | 0.225 | 0.256 | 0.069 |

**Table S4:**
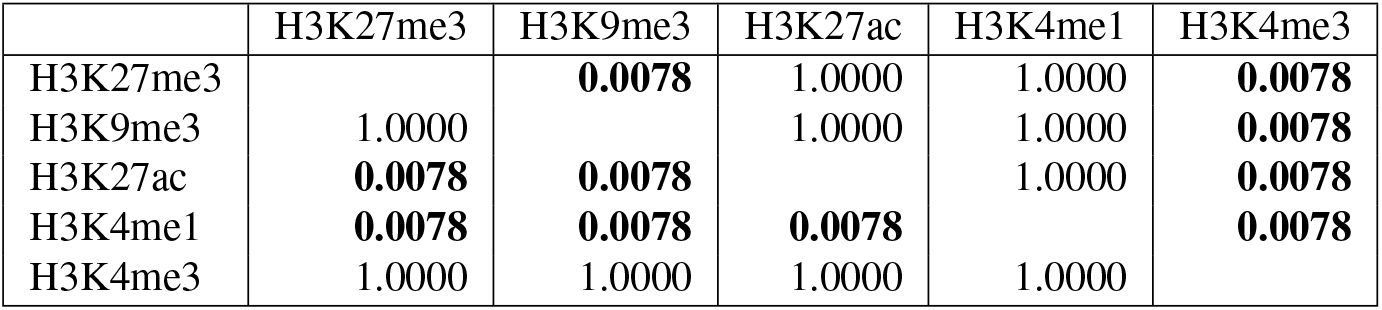
p-values for paired Wilcoxon one-sided (line greater than column) test between the different marks on the mouse brain data.

|  | H3K27me3 | H3K9me3 | H3K27ac | H3K4me1 | H3K4me3 |
| --- | --- | --- | --- | --- | --- |
| H3K27me3 |  | **0.0078** | 1.0000 | 1.0000 | **0.0078** |
| H3K9me3 | 1.0000 |  | 1.0000 | 1.0000 | **0.0078** |
| H3K27ac | **0.0078** | **0.0078** |  | 1.0000 | **0.0078** |
| H3K4me1 | **0.0078** | **0.0078** | **0.0078** |  | **0.0078** |
| H3K4me3 | 1.0000 | 1.0000 | 1.0000 | 1.0000 |  |

**Table S5:**
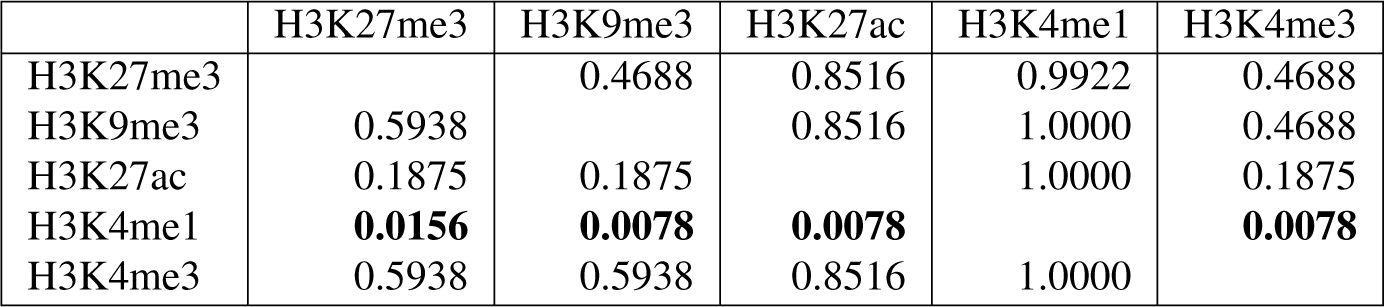
p-values for paired Wilcoxon one-sided (line greater than column) test between the different marks on the human PBMC data.

**Table S6:** p-values for paired Wilcoxon one-sided (line greater than column) test between the different methods accross both the human PBMC and mouse brain data.

|  | Signac | ChromScape_LSI | PeakVI | SnapATAC | cisTopic | SCALE | ChromScape_PCA |
| --- | --- | --- | --- | --- | --- | --- | --- |
| Signac |  | 0.9863 | 0.0801 | 0.1377 | 0.0801 | **0.0322** | **0.0010** |
| ChromScape_LSI | **0.0186** |  | **0.0020** | **0.0049** | **0.0029** | **0.0020** | **0.0010** |
| PeakVI | 0.9346 | 0.9990 |  | 0.7842 | 0.3125 | **0.0098** | **0.0010** |
| SnapATAC | 0.8838 | 0.9971 | 0.2461 |  | 0.0801 | 0.0527 | **0.0049** |
| cisTopic | 0.9346 | 0.9980 | 0.7217 | 0.9346 |  | **0.0068** | **0.0010** |
| SCALE | 0.9756 | 0.9990 | 0.9932 | 0.9580 | 0.9951 |  | **0.0010** |
| ChromScape_PCA | 1.0000 | 1.0000 | 1.0000 | 0.9971 | 1.0000 | 1.0000 |  |

**Table S7:**
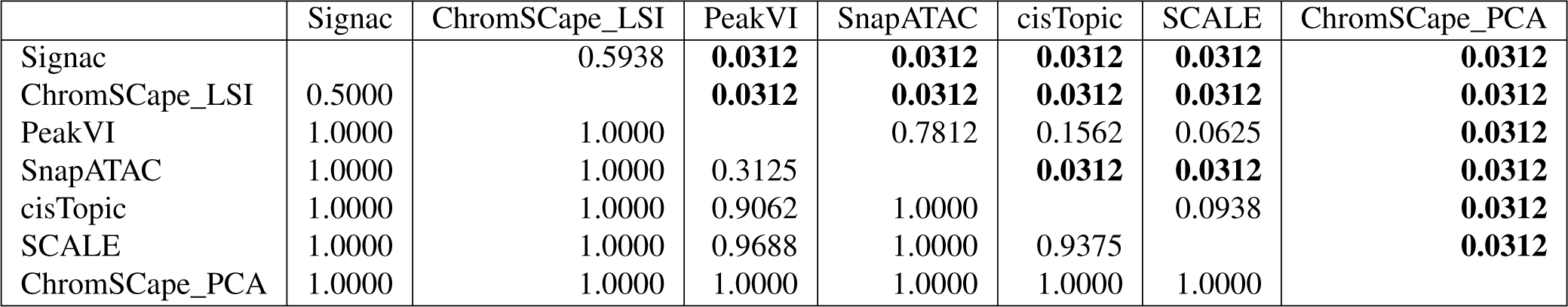
p-values for paired Wilcoxon one-sided (line greater than column) test between the different methods on the mouse brain data.

|  | Signac | ChromScape_LSI | PeakVI | SnapATAC | cisTopic | SCALE | ChromScape_PCA |
| --- | --- | --- | --- | --- | --- | --- | --- |
| Signac |  | 0.5938 | **0.0312** | **0.0312** | **0.0312** | **0.0312** | **0.0312** |
| ChromScape_LSI | 0.5000 |  | **0.0312** | **0.0312** | **0.0312** | **0.0312** | **0.0312** |
| PeakVI | 1.0000 | 1.0000 |  | 0.7812 | 0.1562 | 0.0625 | **0.0312** |
| SnapATAC | 1.0000 | 1.0000 | 0.3125 |  | **0.0312** | **0.0312** | **0.0312** |
| cisTopic | 1.0000 | 1.0000 | 0.9062 | 1.0000 |  | 0.0938 | **0.0312** |
| SCALE | 1.0000 | 1.0000 | 0.9688 | 1.0000 | 0.9375 |  | **0.0312** |
| ChromScape_PCA | 1.0000 | 1.0000 | 1.0000 | 1.0000 | 1.0000 | 1.0000 |  |

**Table S8:**
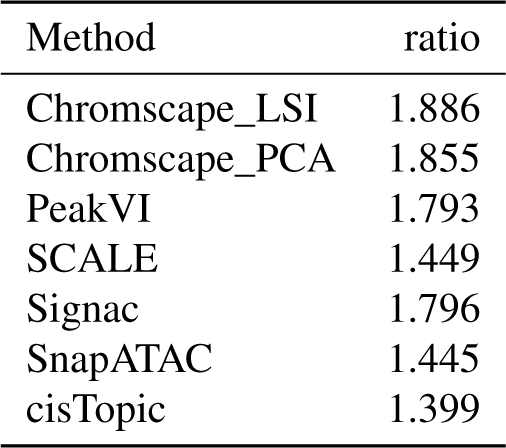
Ratio between the best and worst performances for each method across matrix construction, with no preprocessing, averaged over the mouse brain dataset marks.

**Table S9:** Ratio between the best and worst performances for each mark of the mouse brain dataset across matrix construction, with no preprocessing, averaged over the 7 methods.

| Mark | ratio |
| --- | --- |
| H3K27ac | 1.517 |
| H3K27me3 | 2.022 |
| H3K4me1 | 1.550 |
| H3K4me3 | 1.606 |
| H3K9me3 | 1.608 |

**Table S10:** Ratio between the best and worst performances across matrix construction on the raw data (no feature or cell selection applied) for each mark and method combination on the mouse brain dataset.

|  | H3K27me3 | H3K9me3 | H3K27ac | H3K4me1 | H3K4me3 |
| --- | --- | --- | --- | --- | --- |
| Signac | 1.96 | 1.96 | 1.65 | 1.79 | 1.62 |
| ChromSCape_LSI | 2.54 | 1.95 | 1.37 | 1.22 | 2.35 |
| PeakVI | 2.83 | 1.57 | 1.65 | 1.36 | 1.57 |
| SnapATAC | 1.58 | 1.61 | 1.28 | 1.37 | 1.38 |
| cisTopic | 1.43 | 1.35 | 1.35 | 1.54 | 1.33 |
| SCALE | 1.80 | 1.33 | 1.38 | 1.43 | 1.30 |
| ChromSCape_PCA | 2.01 | 1.49 | 1.94 | 2.14 | 1.70 |

**Table S11:** Ratio between the best and worst performances across matrix construction on the raw data (no feature or cell selection applied) for each mark and method combination on the human PBMC dataset.

|  | H3K27me3 | H3K9me3 | H3K27ac | H3K4me1 | H3K4me3 |
| --- | --- | --- | --- | --- | --- |
| Signac | 2.78 | 5.16 | 4.41 | 2.47 | 1.18 |
| ChromSCape_LSI | 1.30 | 1.56 | 1.52 | 1.25 | 1.46 |
| PeakVI | 1.20 | 3.95 | 5.07 | 7.64 | 1.10 |
| SnapATAC | 2.65 | 2.36 | 1.00 | 3.19 | 2.57 |
| cisTopic | 1.04 | 1.12 | 1.18 | 1.19 | 1.40 |
| SCALE | 2.00 | 2.14 | 2.22 | 3.19 | 1.12 |
| ChromSCape_PCA | 1.14 | 1.32 | 1.48 | 1.27 | 1.07 |

**Table S12:** Ratio of the performances between the best coverage threshold and the worst one for each mark on the mouse brain dataset.

| Mark | best increase Signac | best increase ChromSCape_LSI |
| --- | --- | --- |
| H3K27me3 | 1.104 | 1.071 |
| H3K9me3 | 1.073 | 1.043 |
| H3K27ac | 1.110 | 1.095 |
| H3K4me1 | 1.149 | 1.133 |
| H3K4me3 | 1.008 | 1.082 |

**Table S13:** Ratio of the performances between between the best coverage threshold and the worst one, averaged by method on the mouse brain dataset.

| Method | best increase |
| --- | --- |
| Chromscape_LSI | 1.085 |
| Chromscape_PCA | 1.409 |
| PeakVI | 1.076 |
| SCALE | 1.211 |
| Signac | 1.089 |
| SnapATAC | 1.098 |
| cisTopic | 1.082 |

**Table S14:** Ratio of the performances, averaged by mark, between having all the cells present and having either only 40% or 60% of themin the mouse brain dataset.

| Mark | Signac |  | ChromScape_LSI |  |
| --- | --- | --- | --- | --- |
|  | From 40% | From 60% | From 40% | From 60% |
| H3K27ac | 1.091 | 1.047 | 1.088 | 1.028 |
| H3K27me3 | 1.155 | 1.061 | 1.118 | 1.084 |
| H3K9me3 | 1.143 | 1.089 | 1.096 | 1.017 |
| H3K4me1 | 1.057 | 1.013 | 1.057 | 1.025 |
| H3K4me3 | 1.172 | 1.081 | 1.185 | 1.075 |

**Table S15:** Ratio of the performances, averaged by method, between having all the cells present and having either only 40% or 60% of themin the mouse brain dataset.

| Method | From 40% | From 60% |
| --- | --- | --- |
| Chromscape_LSI | 1.109 | 1.046 |
| Chromscape_PCA | 1.188 | 1.092 |
| PeakVI | 1.342 | 1.180 |
| SCALE | 1.290 | 1.159 |
| Signac | 1.124 | 1.058 |
| SnapATAC | 1.084 | 1.038 |
| cisTopic | 1.210 | 1.108 |

**Table S16:** Ratio of the performances between having high coverage (q50_100 condition) in the mouse brain dataset, and either low coverage or baseline coverage, averaged by mark. The “large increase” column is the increase in performance observed against q0_50 where we select the cells with the lowest coverage. The “baseline” column is the increase against no selection.

| Mark | Signac |  | ChromScape_LSI |  |
| --- | --- | --- | --- | --- |
|  | large increase | baseline | large increase | baseline |
| H3K27ac | 1.708 | 1.150 | 1.666 | 1.119 |
| H3K27me3 | 1.711 | 1.216 | 1.464 | 1.170 |
| H3K9me3 | 1.555 | 1.223 | 1.355 | 1.140 |
| H3K4me1 | 1.688 | 1.193 | 1.754 | 1.204 |
| H3K4me3 | 1.682 | 1.087 | 2.067 | 1.220 |

**Table S17:** Ratio of the performances between having high coverage (q50_100 condition) in the mouse brain dataset, and either low coverage or baseline coverage, averaged by method. The “large increase” column is the increase in performance observed against q0_50 where we select the cells with the lowest coverage. The “baseline” column is the increase against no selection.

| Method | large increase | baseline |
| --- | --- | --- |
| Chromscape_LSI | 1.661 | 1.170 |
| Chromscape_PCA | 1.791 | 1.502 |
| PeakVI | 1.561 | 1.152 |
| SCALE | 1.504 | 1.467 |
| Signac | 1.669 | 1.174 |
| SnapATAC | 1.421 | 1.143 |
| cisTopic | 1.361 | 1.174 |

## Supplementary Figures

**Figure S4:**
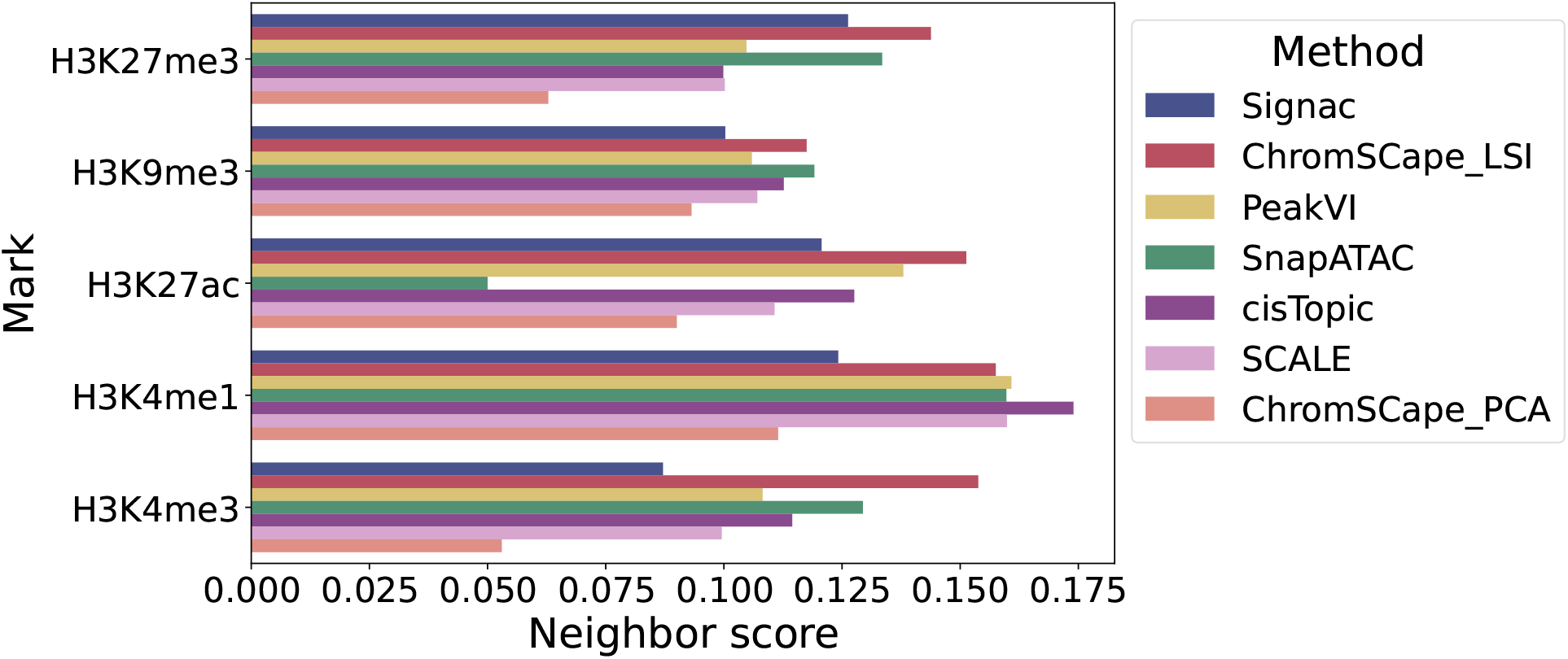
Best performances of the different representation methods on the human PBMC dataset.

**Figure S5:**
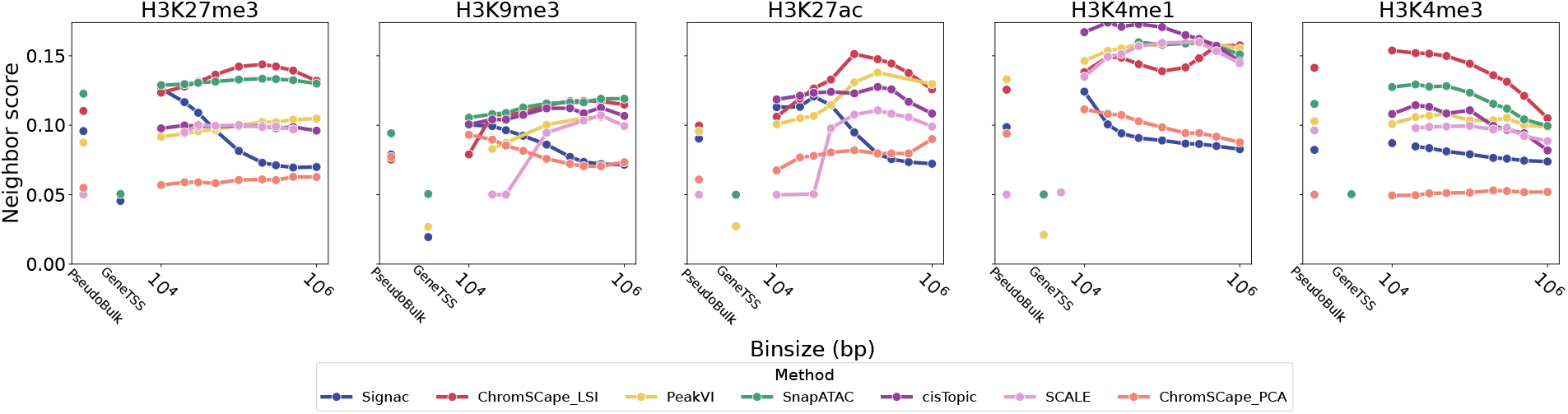
Performances of the 7 dimension reduction algorithms on the 5 marks in the human PBMC dataset, as a function of the matrix construction.

**Figure S6:**
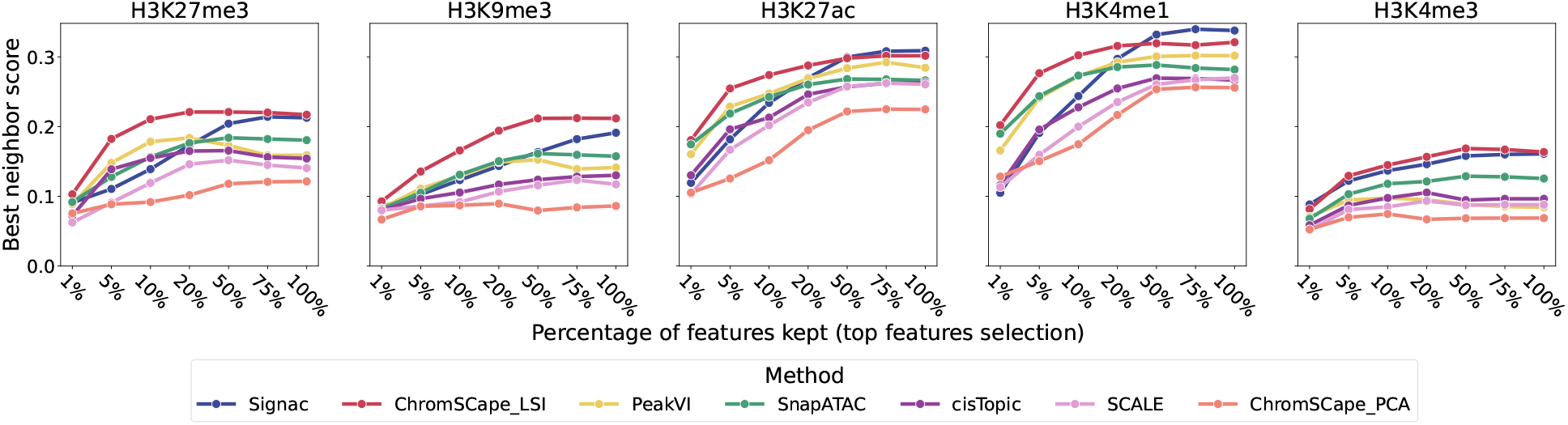
Role of feature selection, using the top features method used for scRNA-seq Each point corresponds to the best performance across matrix construction of a given method and a given percentage of features kept, for the 7 methods, 5 marks, and 7 features selection conditions

## Notes

### Competing Interest Statement

The authors have declared no competing interest.

